# ProteinBERT: A universal deep-learning model of protein sequence and function

**DOI:** 10.1101/2021.05.24.445464

**Authors:** Nadav Brandes, Dan Ofer, Yam Peleg, Nadav Rappoport, Michal Linial

## Abstract

Self-supervised deep language modeling has shown unprecedented success across natural language tasks, and has recently been repurposed to biological sequences. However, existing models and pretraining methods are designed and optimized for text analysis. We introduce ProteinBERT, a deep language model specifically designed for proteins. Our pretraining scheme consists of masked language modeling combined with a novel task of Gene Ontology (GO) annotation prediction. We introduce novel architectural elements that make the model highly efficient and flexible to very large sequence lengths. The architecture of ProteinBERT consists of both local and global representations, allowing end-to-end processing of these types of inputs and outputs. ProteinBERT obtains state-of-the-art performance on multiple benchmarks covering diverse protein properties (including protein structure, post translational modifications and biophysical attributes), despite using a far smaller model than competing deep-learning methods. Overall, ProteinBERT provides an efficient framework for rapidly training protein predictors, even with limited labeled data. Code and pretrained model weights are available at https://github.com/nadavbra/protein_bert.

## Background

Proteins are nature’s ultimate machines, found across the entire tree of life. While knowledge of protein sequences is accumulating exponentially, understanding their functions remains one of the greatest scientific challenges of our time, with numerous implications to human health. Protein sequences can be viewed as strings of amino-acid letters. As such, machine-learning methods developed for natural language and other sequences are a natural fit to predictive protein tasks [1].

Modern deep neural network architectures specifically designed for sequences (such as BERT [2, 3]), combined with pretraining on massive datasets, have led to a revolution in automated text analysis [4]. The attention-based Transformer architecture in particular has shown astounding performance over a wide range of benchmarks across many domains [5, 6].

At the heart of these successes are self-supervised and transfer learning. According to the transfer-learning paradigm, a model is first pre-trained on one task, and then fine-tuned on other downstream tasks of interest [7–9]. Assuming that the pretraining and downstream tasks are somehow related (e.g. both require understanding texts in the same language), pretraining can help the model learn useful representations for the downstream tasks. In self-supervised pretraining, labels are automatically generated, allowing models to learn from enormous, unlabeled datasets [10]. A common example of self-supervised learning is language modeling, where a model (typically a deep neural network) learns language structure by filling missing parts in a text (which have been hidden with a special mask token) or reconstructing corrupted text [11, 12]. Fine-tuning, on the other hand, is typically supervised and requires labeled data. The transfer-learning paradigm has allowed predictive models to achieve substantial performance gains across numerous benchmarks, especially in tasks where labeled data is scarce [13, 14].

Most sequence-based language models (e.g. BERT [2], ULMFiT [11], XLNet [15], ELECTRA [16]) have been designed for processing natural languages (with a bias towards English). Thus, their architectures and pretraining tasks may not be optimal for proteins, which, despite many structural similarities, have different properties from human language [1, 17]. Most notably, proteins do not have clear-cut multi-letter building blocks (such as words and sentences). Moreover, proteins are more variable in length than sentences, and show many interactions between distant positions (due to their 3D structure). To this day, protein research is still dominated by classical sequence-similarity methods (such as BLAST [18] and hidden Markov models [19]), in contrast to domains such as computer vision which have become dominated by deep learning. A few recent studies have pretrained deep neural language models on protein sequences (e.g. ESM [20], TAPE-Transformer [21], UDSMProt [22], UniRep [23]) [24–26]. Such works usually import existing architectures and tasks from the natural language domain, without taking advantage of the unique characteristics of proteins.

Here, we present ProteinBert, a new deep-learning model designed for protein sequences. We improve upon the classic Transformer/BERT architecture, and introduce a novel pretraining task of predicting protein functions. We pretrained ProteinBert on ∼106M proteins (representing the entire known protein space) on two simultaneous tasks. The first task is bidirectional language modeling of protein sequences. The second task is Gene Ontology (GO) annotation prediction, which captures diverse protein functions [27]. GO annotations are a manually curated set of ∼45K terms defined at the whole-protein level, covering the entire protein space across all organisms. They cover molecular functions, biological processes and subcellular locations. Unlike classic Transformers, ProteinBERT separates local (character level) and global (whole protein level) representations, thereby supporting multitasking of both local and global tasks in a principled way. While ProteinBERT is considerably smaller and faster than existing models, it approaches or exceeds state-of-the-art performance on a diverse set of benchmarks.

## Methods

### Data

#### Protein dataset for pretraining

ProteinBERT was pretrained on ∼106M proteins derived from UniProtKB/UniRef90, covering the entire tree of life [28, 29]. UniRef90 provides a non-redundant set of protein clusters sharing at least 90% sequence identity. Each cluster is represented by a single representative protein, ensuring a relatively homogenous coverage of the protein space. For each protein, we extracted its amino-acid sequence and associated GO annotations (according to UniProtKB). We considered only the 8,943 most frequent GO annotations that occurred at least 100 times in UniRef90. Of the ∼106M UniRef90 proteins, ∼46M had at least one of the 8,943 considered annotations (with ∼2.3 annotations per protein, on average across the ∼46M proteins).

#### Protein benchmarks

To evaluate ProteinBERT, we tested it on nine benchmarks concerning all major facets of protein research, covering protein structure, post-translational modifications and biophysical properties (Table 1). Labels in these benchmarks are either local (e.g. post-translational modifications) or global (e.g. remote homology), and they are either continuous (e.g. protein stability), binary (e.g. signal peptide) or categorical (e.g. secondary structure). Notably, in local benchmarks the number of training samples is much greater than the number of protein sequences, as target labels are per residue.

**Table 1:** Protein benchmarks.

| Topic | Benchmark | Target type<sup>a</sup> | Resolution | # Training sequences | Source |
| --- | --- | --- | --- | --- | --- |
| Protein structure | Secondary structure | Categorical (3) | Local | 8,678 | [21, 30] |
|  | Disorder | Binary | Local | 8,678 | [30] |
|  | Remote homology | Categorical (1,195) | Global | 12,312 | [21, 31, 32] |
|  | Fold classes | Categorical (7) | Global | 15,680 | [31, 32] |
| Post- | Signal peptide | Binary | Global | 16,606 | [33] |
| translational modifications | Major PTMs | Binary | Local | 43,356 | [34] |
|  | Neuropeptide cleavage | Binary | Local | 2,727 | [35–37] |
| Biophysical properties | Fluorescence | Continuous | Global | 21,446 | [21, 38] |
|  | Stability | Continuous | Global | 53,679 | [21, 39] |

<sup>a</sup>For categorical targets, the number of classes appears in parentheses.

Four out of nine benchmarks (secondary structure, remote homology, fluorescence and stability) were taken from TAPE (Tasks Assessing Protein Embeddings), a standardized set of benchmarks for evaluating protein sequence models [21]. In addition, we introduce five new benchmarks (see Supplementary Methods).

### Sequence and annotation encoding

Protein sequences were encoded as sequences of integer tokens. We used 26 unique tokens representing the 20 standard amino acids, selenocysteine (*U*), an undefined amino-acid (*X*), another amino acid (*OTHER*), and 3 additional tokens (*START, END* and *PAD*). For each sequence, *START* and *END* tokens were added before the first amino acid and after the last amino acid, respectively. The *PAD* token was added to pad sequences shorter than the sequence length chosen for the minibatch.

The architecture of ProteinBERT (like all deep learning models) dictates that each minibatch has a fixed sequence length. We included the *START* and *END* tokens to help the model interpret proteins that are longer than the chosen sequence length. When encoding a protein longer than the chosen sequence length, we selected a random subsequence of the protein, leaving out at least one of its two ends. The absence of the START or END token allowed the model to recognize that it only received part of a sequence.

GO annotations of every sequence were encoded as a binary vector of fixed size (8,943), where all entries are zeros except those corresponding to GO annotation associated with the protein.

### Self-supervised pretraining on protein sequences and annotations

To learn efficient protein representations, ProteinBERT was pretrained on protein sequences and GO annotations extracted from UniRef90. The model received corrupted inputs (protein sequences and GO annotations) and had to recover the uncorrupted data. The corruption of protein sequences was performed by randomly replacing tokens with 5% probability (i.e. keeping the original token with 95% probability, or replacing it with a uniformly-selected random token with 5% probability). Input GO annotations were corrupted by randomly removing existing annotations with 25% probability, and adding random false annotations with probability of 0.01% for each annotation not associated with the protein. For 50% of the processed proteins, we removed all input annotations altogether, to force the model to predict GO annotations from sequence alone (as was the case in all tested benchmarks). In summary, the described pretraining is a dual task, where the model has to recover both the protein sequence and its known GO annotations. The latter task is relevant to numerous domains of protein research, given the wide range of functions covered by GO terms.

To avoid learning the GO annotations of proteins in the tested benchmarks (Table 1), we ignored the GO annotations of proteins with at least 40% sequence similarity to any record in the test-sets of the benchmarks. To this end, we used BLASTP [40] (with default settings), identifying ∼600K such sequences (out of the ∼106M pretraining proteins).

The loss function minimized by ProteinBERT during pretraining was a sum of the categorical cross-entropy over the protein sequences and the binary cross-entropy over the GO annotations, namely 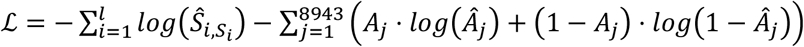, where *l* is the sequence length, *S*_*i*_ ∈ {1, …, 26} is the sequence’s true token at position *i, Ŝ*_*i,k*_ ϵ [0,1] is the predicted probability that position *i* has the token *k* (for any *k* ∈ {1, …, 26}), *A*_*j*_ ∈ {0,1} is the true indicator for annotation *j* (for any *j* ∈ {1, … 8943}), and *Â*_*j*_ ϵ [0,1] is the predicted probability for the protein to have annotation *j*.

An important feature of ProteinBERT is sequence length flexibility. To avoid the risk of overfitting the model to a specific constant length, we periodically (every 15 minutes of training) switched the encoding length of protein sequences, using lengths of 128, 512 or 1,024 tokens.

Pretraining speed on a single GPU (Nvidia Quadro RTX 5000) was 280 protein records per second. We trained the model for 28 days over ∼670M records (i.e. ∼6.4 iterations over the entire training dataset of ∼106M records). The trained model weights are publicly available along with our code (see below).

### Supervised fine-tuning on protein benchmarks

Following pretraining, we fine-tuned and evaluated the model on a diverse set of benchmarks (Table 1). For all benchmarks, ProteinBERT was initialized from the same pretrained state and fine-tuned through the same protocol. Initially, all layers of the pretrained model were frozen, and only a newly added fully-connected layer was allowed to train for up to 40 epochs. Next, we unfroze all the layers and trained the model for up to 40 additional epochs. Finally, we trained the model for 1 final epoch of a larger sequence length (see Supplementary Methods). Throughout all epochs, we reduced the learning rate on plateau and applied early stopping based on an independent validation set. Model evaluation was then performed over a held-out test set. The entire fine-tuning procedure took ∼14 minutes on a single GPU (on average across the nine benchmarks).

### Deep-learning architecture

While inspired by BERT [2], the architecture of ProteinBERT is different and includes several innovations. ProteinBERT is a type of a denoising autoencoder, with corresponding inputs and outputs (Fig. 1). The two inputs (and outputs) of ProteinBERT are i) protein sequences (encoded as token sequences) and ii) protein GO annotations (encoded as fixed-size binary vectors).

**Fig. 1:**
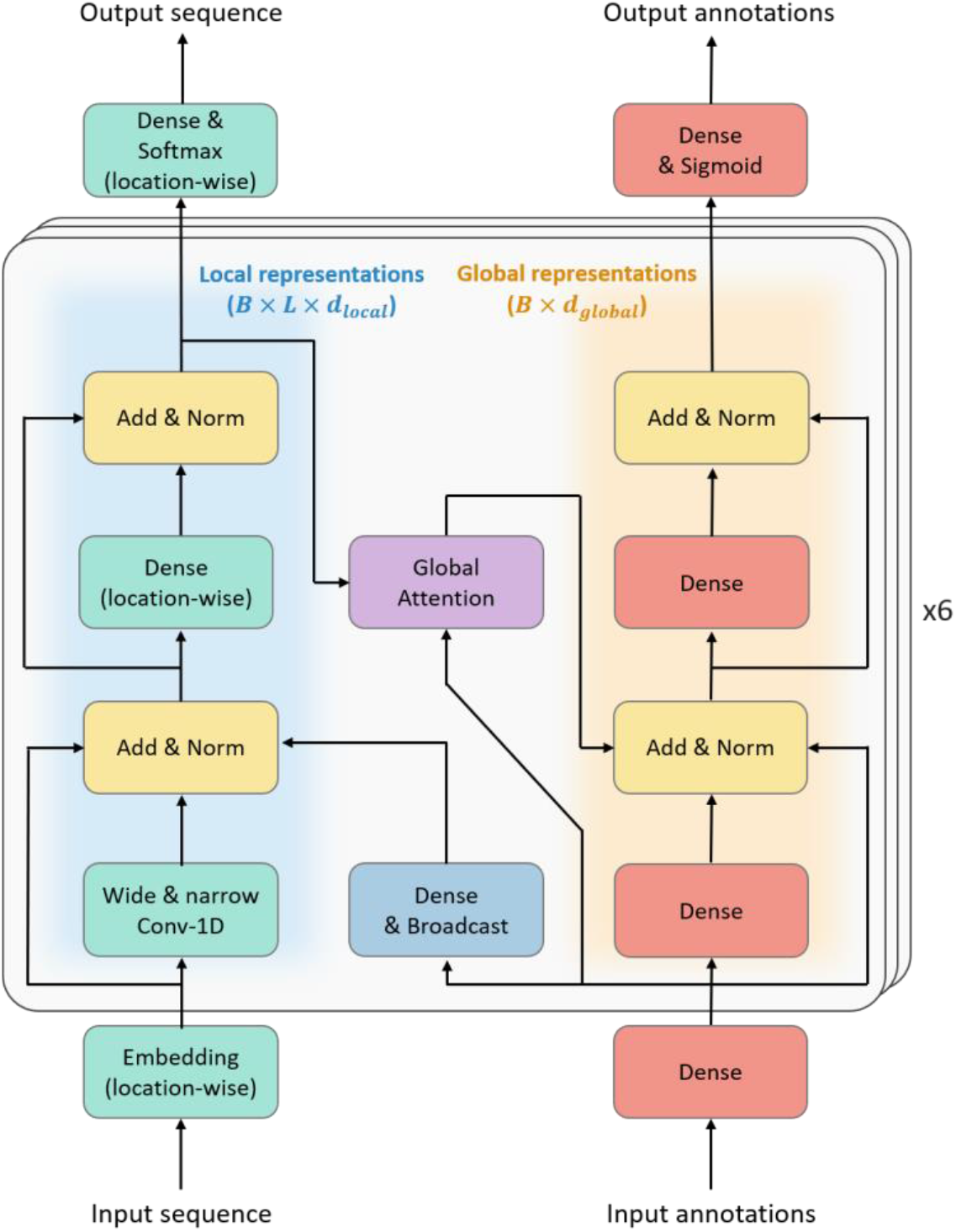
The ProteinBERT architecture. ProteinBERT’s architecture is inspired by BERT. Unlike standard Transformers, ProteinBERT supports both local (sequential) and global data. The model consists of 6 transformer blocks manipulating local (left side) and global (right side) representations. Each such block manipulates these representations by fully-connected and convolutional layers (in the case of local representations), with skip connections and normalization layers between them. The local representations affect the global representations through a global attention layer, and the global representations affect the local representations through a broadcast fully-connected layer.

The architecture of the model consists of two almost parallel paths: one for local representations and the other for global representations (Fig. 1). The local representations are 3D tensors of shape *B* × *L* × *d*_*local*_ where *B* is the batch size, *L* is the minibatch sequence length, and *d*_*local*_ is the number of channels for the local representations (we used *d*_*local*_ = 128). The global representations are 2D tensors of shape *B* × *d*_*global*_ (using *d*_*global*_ = 512). In the first layers of the model, the input sequences are transformed into the local-representation 3D tensor by an embedding layer with *d*_*local*_ output features (which is applied independently and identically position-wise), and the input binary annotations are transformed into the global-representation 2D tensor by a fully-connected layer with *d*_*global*_ output features.

The local and global representations are processed by a series of 6 transformer blocks with skip connections and layer normalizations between their hidden layers. Within each block, the local representation is transformed first by 1D convolutional layers, and then by a (location-wise) fully-connected layer. To allow the local representations at each position to be based on other positions at both short and remote proximity, we used both a narrow (without dilation) and a wide (with dilation rate of 5) convolutional layer. Both types of convolution layers have a kernel size of 9 and stride size of 1. Accordingly, each narrow layer has a receptive field of 9 and each wide layer has a receptive field of 41 over the previous layer, meaning that the 6th transformer block has a receptive field of 241 over the input sequence. The global representations, on the other hand, are transformed by two simple fully-connected layers per block (with normalizations between them). All the hidden fully-connected and convolutional layers of the model use GELU (Gaussian Error Linear Unit) activations [41].

The only information flow between the local and global representations occurs through broadcast fully-connected layers (from the global to the local representations) and global attention layers (from the local to the global representations). The broadcast layers are fully-connected layers that transform the *d*_*global*_ features of the global representation into *d*_*local*_ features of the local representations, and then replicate that representation across each of the *L* sequence positions.

The global attention layer is a novel architectural innovation inspired by self-attention [3]. While self-attention takes an input sequence and outputs another sequence by allowing each position to attend to each other position, global attention takes as input both a sequence and a global fixed-size vector and outputs a global fixed-size vector created by attending to each of the local input positions according to the global input vector. Formally, a single-head global attention layer takes as inputs a global representation vector 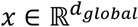 and local representation vectors across *L* positions, 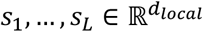, and outputs a global output 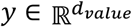. Similar to self-attention, the output is calculated by 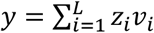 where 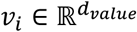 is the value associated with each position *i* ∈ {1, … *L}* and z_i_ ∈ [0,1] is the amount of attention allocated to that position (satisfying z_1_ + ⋯ + z_*L*_ = 1). Like in self-attention, the value associated with each position is calculated by *v*_*i*_ = *σ*(*W*_*v*_*s*_*i*_), using a parameter matrix 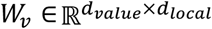 and an activation function *σ* (we chose GELU). Attention values are calculated by 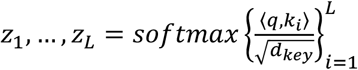, based on query and key vectors 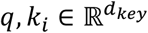. Notice that while the key vectors *k*_1_, …, *k*_*L*_ are specific to each position, the query vector *q* is global. Like in self-attention, the key vectors are calculated by *k*_*i*_ = tanh(*W*_*k*_*s*_*i*_), using a second parameter matrix 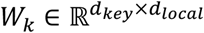. The global query vector is calculated by *q* = tanh(*W*_*q*_*x*), using a third parameter matrix 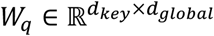. Overall, a single-head global attention layer uses three parameter matrices fit during training, *W*_*q*_, *W*_*k*_ and *W*_*v*_. It is also parameterized by the key dimension *d*_*key*_ (we used *d*_*key*_ = 64). A multi-head global attention layer is obtained by applying *n*_*heads*_ independent single-head global attention layers (each with its own parameters) and concatenating their outputs, obtaining an output of dimension *n*_*heads*_ *⋅ d*_*value*_ (we used *n*_*heads*_ = 4 across all 6 transformer blocks of ProteinBERT). To satisfy dimensionality constraints, ProteinBERT uses 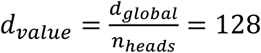

*Overall*, the ProteinBERT model includes 6 transformer blocks with 4 global attention heads in each block. Altogether, it includes ∼16M trainable parameters, making it substantially smaller than other protein language models. For comparison, there are ∼38M parameters in the TAPE Transformer [21], ∼110M in BERT-base [2], and ∼650M in the ESM-1b model [20].

The ProteinBERT architecture has several appealing properties. Most importantly, the entire architecture is agnostic to the length of the processed sequences, and it can be applied over sequences of any given length without changing its learned parameters (our experiments prove that the model indeed generalizes very well across different lengths). This good generality across sequence lengths is also achieved by avoiding positional embeddings used in the standard version of BERT which, in accordance with previous reports [42] and our experimentation, do not always generalize well to sequence lengths longer than those present in the training data. Instead, the convolutional layers and special tokens used at the beginning and end of each sequence provide the model with information on the relative locations of positions. Due to the use of global attention rather than self-attention, the amount of computation performed by the model grows only linearly with sequence length (as opposed to quadratic growth in models with standard self-attention). This linear growth also applies to the model’s memory consumption, allowing ProteinBERT to process extremely long protein sequences (of tens of thousands of amino-acids) intact. Despite this simplification, each position in the local representations and sequence outputs can still depend on the content of each other position, thanks to the alternating information flow between the local and global representations. On top of that, the wide and narrow convolutional layers allow the representation of each position to depend on a large context. By relying on convolutional and attention layers, but avoiding recurrent layers, the computation performed by the network is more efficient and stable with respect to sequence length (as there are no long-term dependencies). Notably, we did not use dropout or any other form of regularization (except for the final fully-connected layer added when fine-tuning the model, which included dropout).

When fine-tuning ProteinBERT on a labeled dataset, another layer is added to its output. The final layer is fed with a concatenation of either the local or global hidden states of the model, depending on whether the output labels are local or global. The activation used for the final layer depends on the output type (i.e. softmax activation for categorical labels, sigmoid activation for binary labels, or no activation for continuous labels).

### Availability

Python code for ProteinBERT’s architecture, pretraining and fine-tuning is open source and available at https://github.com/nadavbra/protein_bert. The repository also includes pretrained model weights and code for downloading and generating the datasets and benchmarks. ProteinBERT is implemented in TensorFlow’s Keras [43, 44].

## Results

### Pretraining improves protein modeling

ProteinBERT was pretrained on ∼106M UniRef90 records for ∼6.4 epochs. We see that the language modeling loss continues to improve on the training set (i.e does not saturate), even after multiple epochs (Fig. 2), in accordance with other studies [20]. The GO annotations task, on the other hand, does show saturation. During pretraining, we periodically changed the sequence length used to encode the input and output protein sequences (128, 512 or 1024 tokens). We observe somewhat lower performance for the 128-token encoding, but similar for 512 and 1024.

**Fig. 2:**
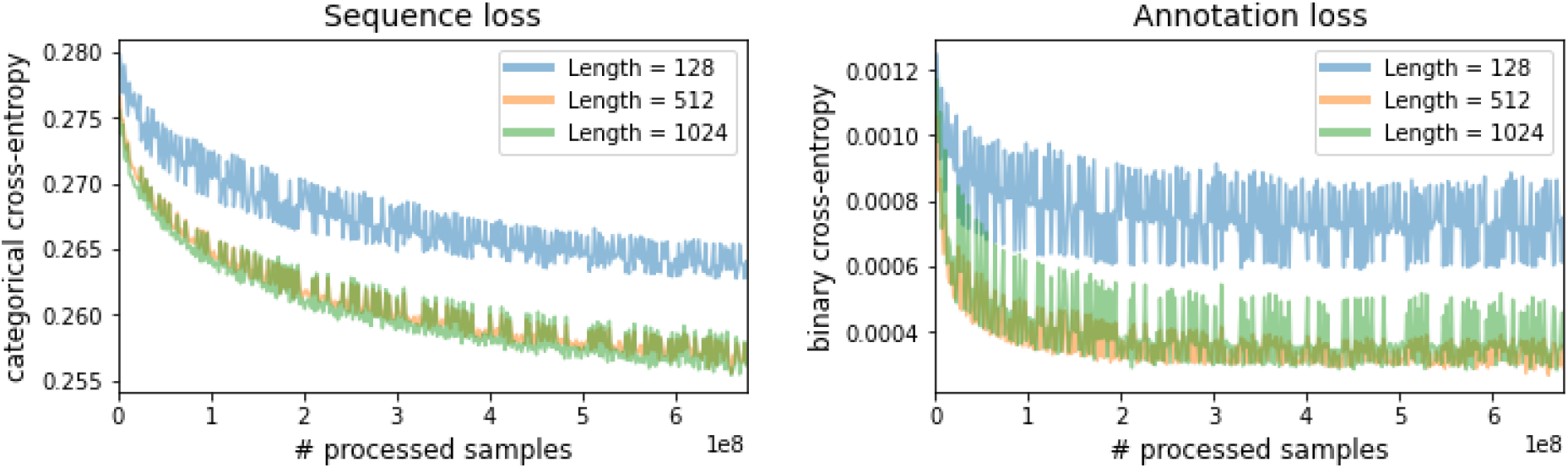
Pretraining loss. Training-set loss over the two pretraining tasks: i) protein sequence language modeling, and ii) GO annotation recovery. Losses were evaluated with input sequence length of 128, 512 or 1,024 tokens on the first 100 batches of the dataset.

### ProteinBERT achieves state-of-the-art results on diverse protein benchmarks

To evaluate ProteinBERT, we used nine benchmarks covering a variety of tasks in protein research (see definitions of the benchmarks in Table 1; full results for all benchmarks are available in Supplementary Table S1). For the four benchmarks taken from TAPE (secondary structure, remote homology, fluorescence and stability prediction), we compared our performance to other state-of-the-art sequence models which had been evaluated on the same benchmarks. Specifically, we compared against a BERT Transformer and LSTM models evaluated in TAPE [21, 23, 45] (Table 2). Notably, the compared deep learning models have ∼38M parameters, in contrast to ∼16M parameters in ProteinBERT. We evaluated ProteinBERT with and without pretraining, observing that pretraining has a major, positive effect on performance in all tasks. Across these benchmarks, ProteinBERT shows performance comparable, or that exceeds similar, larger models, such as the Transformer used in TAPE.

**Table 2:** TAPE benchmark results

|  | Method | Structure | Evolutionary | Engineering |  |
| --- | --- | --- | --- | --- | --- |
|  |  | Secondary structure | Remote homology | Fluorescence | Stability |
| Without Pretraining | TAPE Transformer | 0.70 | 0.09 | 0.22 | -0.06 |
|  | LSTM | 0.71 | 0.12 | 0.21 | 0.28 |
|  | ProteinBERT | 0.70 | 0.06 | 0.65 | 0.63 |
| With Pretraining | TAPE Transformer | 0.73 | 0.21 | 0.68 | 0.73 |
|  | LSTM | 0.75 | 0.26 | 0.67 | 0.69 |
|  | UniRep mLSTM | 0.73 | 0.23 | 0.67 | 0.73 |
|  | ProteinBERT | 0.74 | 0.22 | 0.66 | 0.76 |

To further discern the impact of pretraining on downstream benchmark performance, we evaluated ProteinBERT following different pretraining durations. Specifically, we initiated the model from different snapshots along its pretraining and evaluated its down-stream performance after fine-tuning from these states (Fig. 3). While some tasks do not benefit from pretraining, other tasks (e.g. secondary structure and remote homology) show clear gains from ever more pretraining, and do not show saturation in that improvement. This is notable given that these are among the more challenging tasks.

**Fig. 3:**
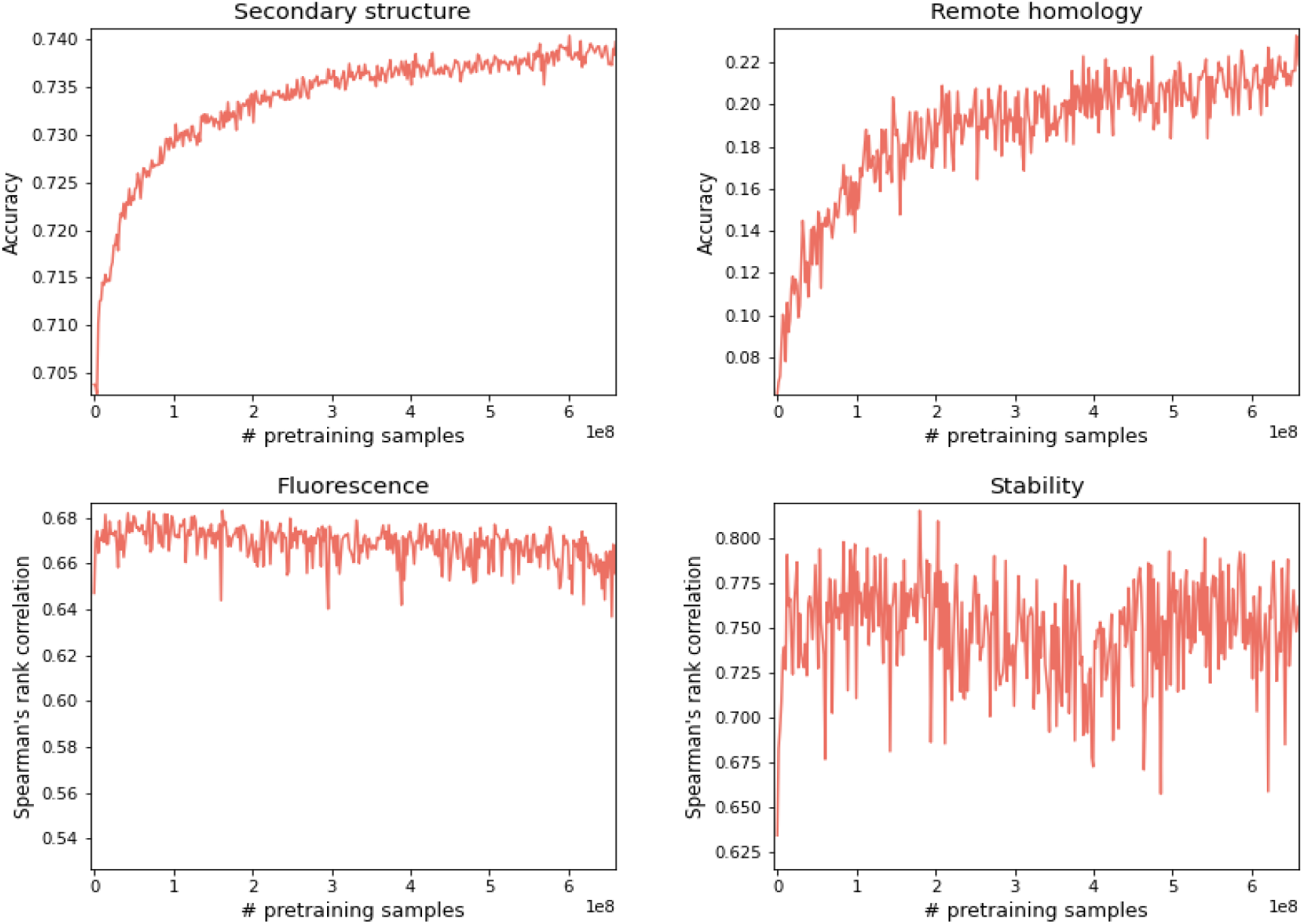
The impact of pretraining on downstream tasks. Performance of fine-tuned ProteinBERT models over the 4 TAPE benchmarks as a function of pretraining amount (measured by the number of processed proteins). Similar plots for all nine benchmarks are shown in Supplementary Fig. S1.

### ProteinBERT generalizes across protein lengths

The architecture of ProteinBERT is efficient and flexible towards different sequence lengths (i.e. the number of tokens encoding the input and output sequences). To test the model’s capacity to generalize across sequence lengths, we measured the test-set performance of ProteinBERT on the 4 of 9 benchmarks that had a non-negligible number of test-set records in proteins longer than 512 tokens (Fig. 4). Specifically, we required at least 25 such records, where a record comprises either an entire protein (in the case of global tasks) or a residue (in the case of local tasks). We observe that in most cases ProteinBERT performs slightly worse for longer sequences, but only modestly, showing that it indeed generalizes across a very wide range of protein lengths. Moreover, the fact that in some cases longer sequences achieve better performance (e.g. 16,384-token sequences in the “Major PTMs” benchmark, or 1,024-token sequences in the “Neuropeptide cleavage” benchmark) suggests that the changes in performance might be due to other factors (e.g. predicting the secondary structure of longer sequences might be an inherently more difficult task).

**Fig. 4:**
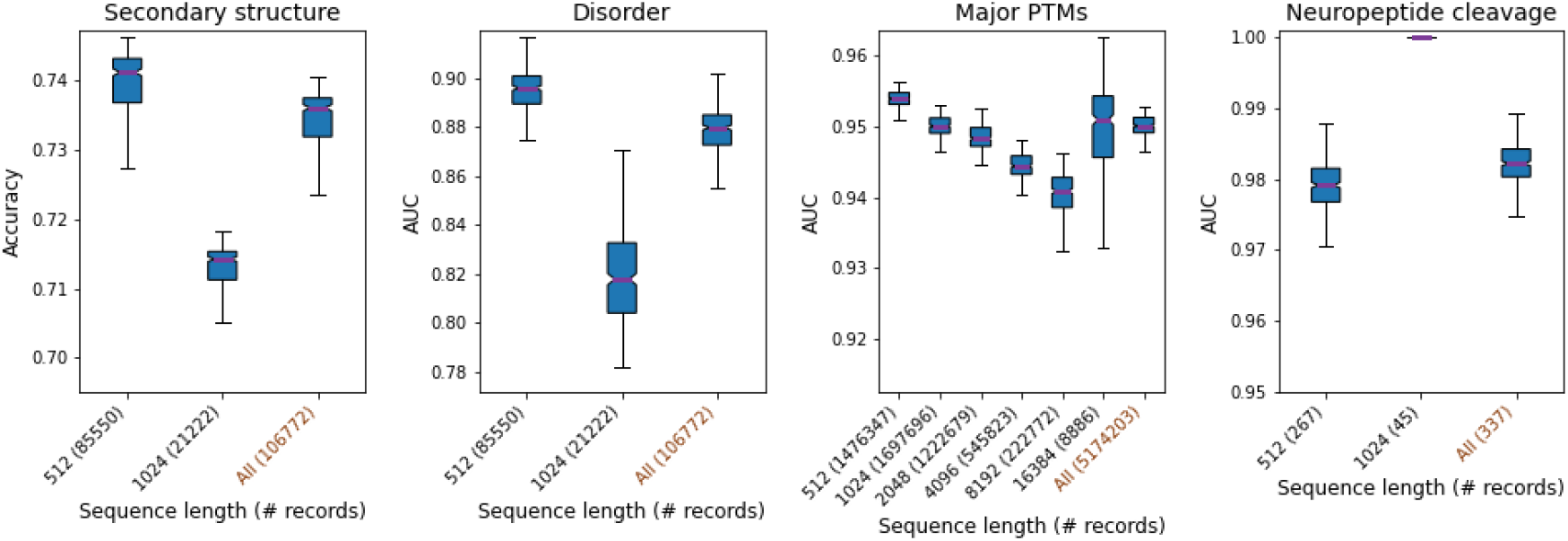
Performance across sequence lengths. Test-set performance of fine-tuned ProteinBERT models with different input sequence lengths. Sequence lengths (e.g. 512, 1,024, etc.) always encode proteins of shorter lengths (e.g. a protein of 700 residues will be encoded as a 1,024-long sequence). Boxplot distributions are over the 371 pretraining snapshots used in Fig. 3.

### Understanding global attention

To demonstrate the inner workings of the global attention mechanism, we extracted the values of the 24 attention heads in ProteinBERT for two proteins selected from the test-set of the signal peptide benchmark, before and after fine-tuning the model on that task (Fig. 5). The patterns of global attention are clearly distinct across different proteins, but some shared patterns exist. For example, attention head #3 in the 3rd transformer block tends to concentrate on the beginning of protein sequences, while attention head #2 in the same layer tends to concentrate on the other parts (i.e. the middle and end of sequences). Fine-tuning the model on signal peptide prediction appears to have mostly altered the last (6th) global attention layer. For example, attention head #1 in that layer changed to emphasize more the beginning of sequences. It is worth stressing that the exact attention values are dependent on the model weights obtained from training, which can change between runs. From our experience, fine-tuning tends to produce rather consistent results, but small differences are sometimes observed.

**Fig. 5:**
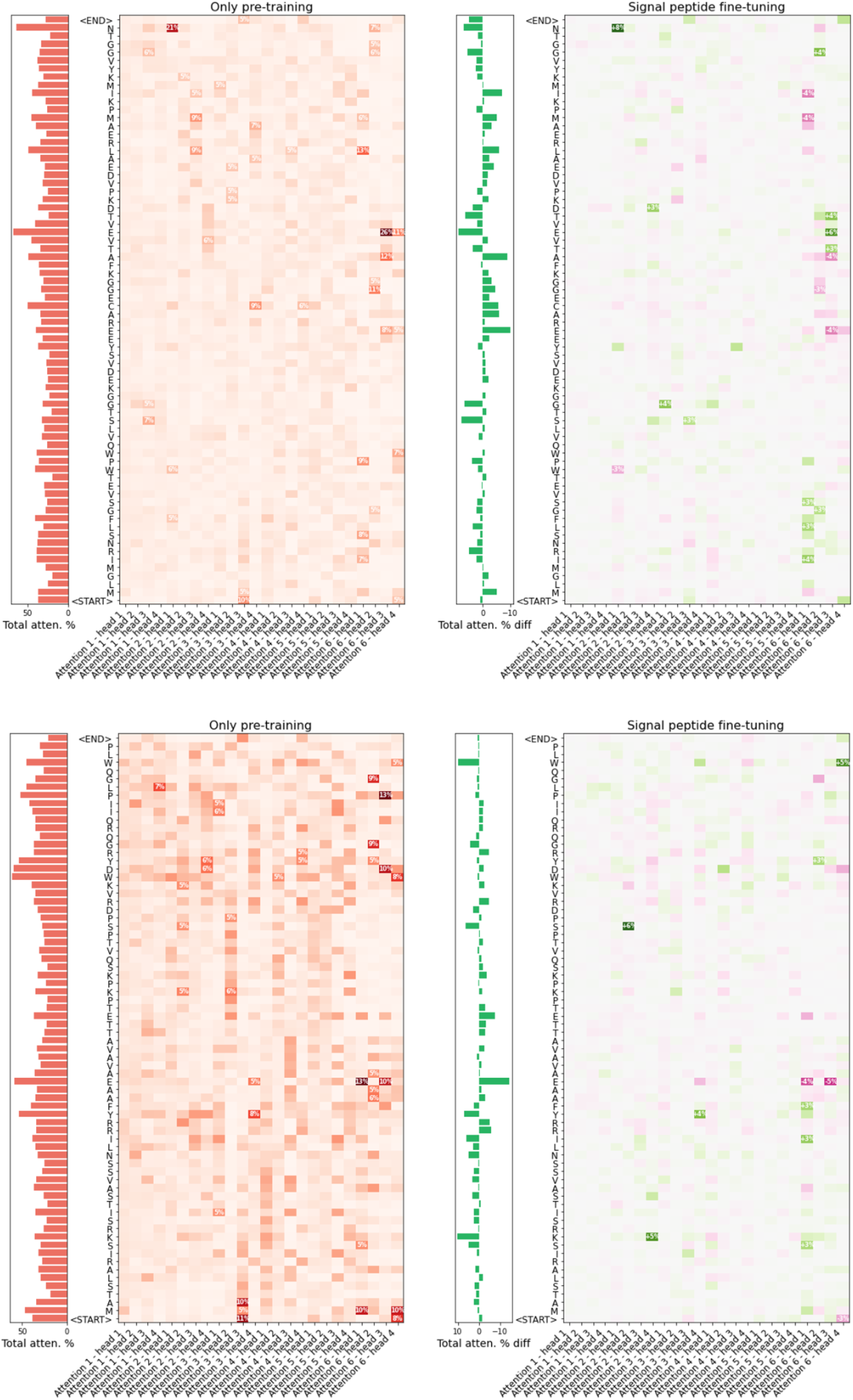
Global attention before and after fine-tuning on signal peptide prediction. Global attention values obtained for two selected proteins: Heme-binding protein 1 (Hebp1) in mouse (top), and Gamma carbonic anhydrase-like 2, mitochondrial protein (GAMMACAL2) in arabidopsis (bottom). The left panels (red colors) show the attention values obtained by the generic ProteinBERT model, after pretraining it as a language model on UniRef90 (but before fine-tuning it on any specific task). The heatmap shows the global attention values at each residue of the protein by each of the 24 attention heads of the model. The bar plot shows the total attention at each residue by summing the attention values across all attention heads. The right panels show the difference in attention values after fine-tuning ProteinBERT on the signal peptide prediction benchmark. The heatmap shows the increase (green) or decrease (purple) of attention across all positions and attention heads. The bar plot shows the total difference in attention at each residue by summing the differences across all attention heads. Notice that each attention head necessarily sums up to 100%. Accordingly, differences sum up to 0%.

## Discussion

We have presented ProteinBERT, a new deep language model for protein sequences designed to capture local and global representations of proteins in a natural way (Fig. 1). We have demonstrated the universality of the model, showing that it can be fine-tuned on a wide range of protein tasks in a matter of minutes and achieve state-of-the-art results (Table 2).

To pretrain ProteinBERT, we introduce a novel pretraining task of protein annotation prediction which is highly suited to proteins (unlike sentence order prediction and other natural language centric tasks [46]). We argue that GO annotations [27] are a sensible extension to language modeling in proteins. They are ubiquitous and available for a large portion of curated proteins (∼46M of the ∼106M proteins in the UniRef90 dataset). Additionally, they can teach the model about a wide range of protein functions (from subcellular localizations to pathways to biochemical roles).

Unlike previous works which included ∼250M putative, redundant sequences [20], we constrained the pretraining of ProteinBERT to ∼106M representative proteins taken from UniRef90 [29], out of the entire known protein space of ∼215M proteins in UniProt [28]. We argue that using a non-redundant set of proteins is more sensible and eliminates a lot of unnecessary bias caused by uneven sampling of the protein space, which is prevalent in the non-filtered version of UniProt. For example, there are >1M proteins in UniProt from the proteome of human immunodeficiency virus 1 (HIV-1), even though the real virus contains only 9 proteins. Such redundancy reflects the abundance of sequence variations along HIV-1 evolution, and the great interest that researchers have had in this variation (compared to most other, far less studied organisms). Using a non-redundant set of proteins is also more efficient, especially when pretraining the model for less than an entire epoch (such as when searching for optimal hyper-parameter combinations).

ProteinBERT’s architecture is efficient and highly scalable, allowing it to process protein sequences of any length. The same model weights conform to any sequence length, allowing it to be trained on a specific range of lengths and then generalize to other, unseen sequence lengths (Fig. 4). By supporting extremely long sequences (more than tens of thousands of residues), ProteinBERT spares the complication of splitting long sequences into smaller chunks, a common practice with self-attention-based models which grow quadratically (rather than linearly) with sequence length [47, 48]. At the core of the model’s flexibility is its use of global attention layers, a new architectural innovation. The compactness of global attention (relative to self-attention) also allows easier inspection of the model’s attention, as all attention values (across all positions and attention heads) can be displayed as a simple 2D map (Fig. 5), as opposed to the 3D map that would be required to cover all-by-all self-attention.

Compatible with the general trends in the field of language modeling [5], we observe that longer pretraining of ProteinBERT shows clear performance gains, both as a language model (Fig. 2) and across many specific tasks (Fig. 3, Supplementary Fig. S1). Existing works show that, other things being equal, larger models and additional pretraining computation time correlates with improved model performance [2, 5, 20]. Thus, we expect larger versions of ProteinBERT (e.g. with more, wider layers) to yield additional improvements. Yet, even with the modest computing resources used in this work (a single GPU), ProteinBERT competes with state-of-the-art models (Table 2), providing a simple and efficient out-of-the-box solution for a wide range of protein tasks. The representations learned by the model through its pretraining are universally applicable across a wide array of tasks, making it useful for few-shot-learning tasks involving limited labelled data.

To facilitate easy usage of ProteinBERT, we provide the pretrained model as a Python package (based on TensorFlow and Keras [43, 44]), which allows automatic downloading of a pretrained model state, fine-tuning and evaluation on labeled datasets.

By providing an effective and accessible model of protein sequence and function, we hope to expedite the adoption of deep language modeling by the protein research community and allow this new powerful tool to further push the boundaries of protein research.

## Author contribution

Conception and study design: Dan, Nadav B, Michal; data acquisition: Nadav B, Dan, Nadav R; architecture design: Nadav B, Dan, Yam; analysis: Nadav B; interpretation of results: Dan, Nadav B, Michal; open sourcing: Nadav B; drafting the manuscript: Nadav B, Dan, Michal.

## Supplementary materials

### Supplementary Methods

**Supplementary Table S1 - Full benchmark results**: Test-set performance of ProteinBERT (following fine-tuning) over all nine benchmarks across 371 snapshots along the pretraining process.

**Supplementary Figure S1**

## References

1. Ofer D, Brandes N, Linial M (2021) The language of proteins: NLP, machine learning \& protein sequences. Comput Struct Biotechnol J

2. Devlin J, Chang M-W, Lee K, Toutanova K (2018) Bert: Pre-training of deep bidirectional transformers for language understanding. arXiv Prepr arXiv 181004805

3. Vaswani A, Shazeer N, Parmar N, et al (2017) Attention is all you need. arXiv Prepr arXiv 170603762

4. Radford A, Wu J, Child R, et al (2019) Language models are unsupervised multitask learners. OpenAI blog 1:9

5. Brown TB, Mann B, Ryder N, et al (2020) Language models are few-shot learners. arXiv Prepr arXiv 200514165

6. Keskar NS, McCann B, Varshney LR, et al (2019) Ctrl: A conditional transformer language model for controllable generation. arXiv Prepr arXiv 190905858

7. Raffel C, Shazeer N, Roberts A, et al (2019) Exploring the limits of transfer learning with a unified text-to-text transformer. arXiv Prepr arXiv 191010683

8. Do CB, Ng AY (2005) Transfer learning for text classification. Adv Neural Inf Process Syst 18:299–306

9. Pan SJ, Yang Q (2009) A survey on transfer learning. IEEE Trans Knowl Data Eng 22:1345–1359

10. Chen T, Kornblith S, Swersky K, et al (2020) Big self-supervised models are strong semi-supervised learners. arXiv Prepr arXiv 200610029

11. Howard J, Ruder S (2018) Universal language model fine-tuning for text classification. arXiv Prepr arXiv 180106146

12. Radford A, Narasimhan K, Salimans T, Sutskever I (2018) Improving language understanding by generative pre-training

13. Thrun S (1996) Is learning the n-th thing any easier than learning the first? In: Advances in neural information processing systems. pp 640–646

14. Wang A, Pruksachatkun Y, Nangia N, et al (2019) Superglue: A stickier benchmark for general-purpose language understanding systems. arXiv Prepr arXiv 190500537

15. Yang Z, Dai Z, Yang Y, et al (2019) Xlnet: Generalized autoregressive pretraining for language understanding. arXiv Prepr arXiv 190608237

16. Clark K, Luong M-T, Le Q V, Manning CD (2020) Electra: Pre-training text encoders as discriminators rather than generators. arXiv Prepr arXiv 200310555

17. Strait BJ, Dewey TG (1996) The Shannon information entropy of protein sequences. Biophys J 71:148–155

18. Altschul SF, Gish W, Miller W, et al (1990) Basic local alignment search tool. J Mol Biol 215:403–410

19. Finn RD, Bateman A, Clements J, et al (2014) Pfam: the protein families database. Nucleic Acids Res 42:D222–30. https://doi.org/10.1093/nar/gkt1223

20. Rives A, Meier J, Sercu T, et al (2021) Biological structure and function emerge from scaling unsupervised learning to 250 million protein sequences. Proc Natl Acad Sci 118:

21. Rao R, Bhattacharya N, Thomas N, et al (2019) Evaluating protein transfer learning with tape. Adv Neural Inf Process Syst 32:9689

22. Strodthoff N, Wagner P, Wenzel M, Samek W (2020) UDSMProt: universal deep sequence models for protein classification. Bioinformatics 36:2401–2409

23. Alley EC, Khimulya G, Biswas S, et al (2019) Unified rational protein engineering with sequence-based deep representation learning. Nat Methods 16:1315–1322

24. Heinzinger M, Elnaggar A, Wang Y, et al (2019) Modeling aspects of the language of life through transfer-learning protein sequences. BMC Bioinformatics 20:1–17

25. Madani A, McCann B, Naik N, et al (2020) Progen: Language modeling for protein generation. arXiv Prepr arXiv 200403497

26. Nambiar A, Heflin M, Liu S, et al (2020) Transforming the language of life: Transformer neural networks for protein prediction tasks. In: Proceedings of the 11th ACM International Conference on Bioinformatics, Computational Biology and Health Informatics. pp 1–8

27. Ashburner M, Ball CA, Blake JA, et al (2000) Gene ontology: tool for the unification of biology. Nat Genet 25:25–29

28. Boutet E, Lieberherr D, Tognolli M, et al (2016) UniProtKB/Swiss-Prot, the manually annotated section of the UniProt KnowledgeBase: how to use the entry view. In: Plant Bioinformatics. Springer, pp 23–54

29. Suzek BE, Huang H, McGarvey P, et al (2007) UniRef: comprehensive and non-redundant UniProt reference clusters. Bioinformatics 23:1282–1288

30. Moult J, Fidelis K, Kryshtafovych A, et al (2018) Critical assessment of methods of protein structure prediction (CASP)—Round XII. Proteins Struct Funct Bioinforma 86:7–15

31. Andreeva A, Howorth D, Chothia C, et al (2014) SCOP2 prototype: a new approach to protein structure mining. Nucleic Acids Res 42:D310–4. https://doi.org/10.1093/nar/gkt1242

32. Andreeva A, Kulesha E, Gough J, Murzin AG (2020) The SCOP database in 2020: expanded classification of representative family and superfamily domains of known protein structures. Nucleic Acids Res 48:D376-D382

33. Armenteros JJA, Tsirigos KD, Sønderby CK, et al (2019) SignalP 5.0 improves signal peptide predictions using deep neural networks. Nat Biotechnol 37:420–423

34. Hornbeck P V, Zhang B, Murray B, et al (2015) PhosphoSitePlus, 2014: mutations, PTMs and recalibrations. Nucleic Acids Res 43:D512--D520

35. Ofer D, Linial M (2015) ProFET: Feature engineering captures high-level protein functions. Bioinformatics 31:3429–3436

36. Brandes N, Ofer D, Linial M (2016) ASAP: A machine learning framework for local protein properties. Database 2016:. https://doi.org/10.1093/database/baw133

37. Ofer D, Linial M (2014) NeuroPID: a predictor for identifying neuropeptide precursors from metazoan proteomes. Bioinformatics 30:931–940

38. Sarkisyan KS, Bolotin DA, Meer M V, et al (2016) Local fitness landscape of the green fluorescent protein. Nature 533:397–401

39. Rocklin GJ, Chidyausiku TM, Goreshnik I, et al (2017) Global analysis of protein folding using massively parallel design, synthesis, and testing. Science (80-) 357:168–175

40. Altschul SF, Madden TL, Schäffer AA, et al (1997) Gapped BLAST and PSI-BLAST : a new generation of protein database search programs. 25:3389–3402

41. Hendrycks D, Gimpel K (2016) Gaussian error linear units (gelus). arXiv Prepr arXiv 160608415

42. Neishi M, Yoshinaga N (2019) On the relation between position information and sentence length in neural machine translation. In: Proceedings of the 23rd Conference on Computational Natural Language Learning (CoNLL). pp 328–338

43. Abadi M, Barham P, Chen J, et al (2016) Tensorflow: A system for large-scale machine learning. In: 12th $\{$USENIX$\}$ symposium on operating systems design and implementation ($\{$OSDI$\}$ 16). pp 265–283

44. Chollet F, others (2015) keras

45. Bepler T, Berger B (2019) Learning protein sequence embeddings using information from structure. arXiv Prepr arXiv 190208661

46. Lan Z, Chen M, Goodman S, et al (2019) Albert: A lite bert for self-supervised learning of language representations. arXiv Prepr arXiv 190911942

47. Choromanski K, Likhosherstov V, Dohan D, et al (2020) Rethinking attention with performers. arXiv Prepr arXiv 200914794

48. Zaheer M, Guruganesh G, Dubey A, et al (2020) Big bird: Transformers for longer sequences. arXiv Prepr arXiv 200714062

