## Supplementary Materials for "ProteinBERT: A universal deep-learning model of protein sequence and function"

### Supplementary Figures

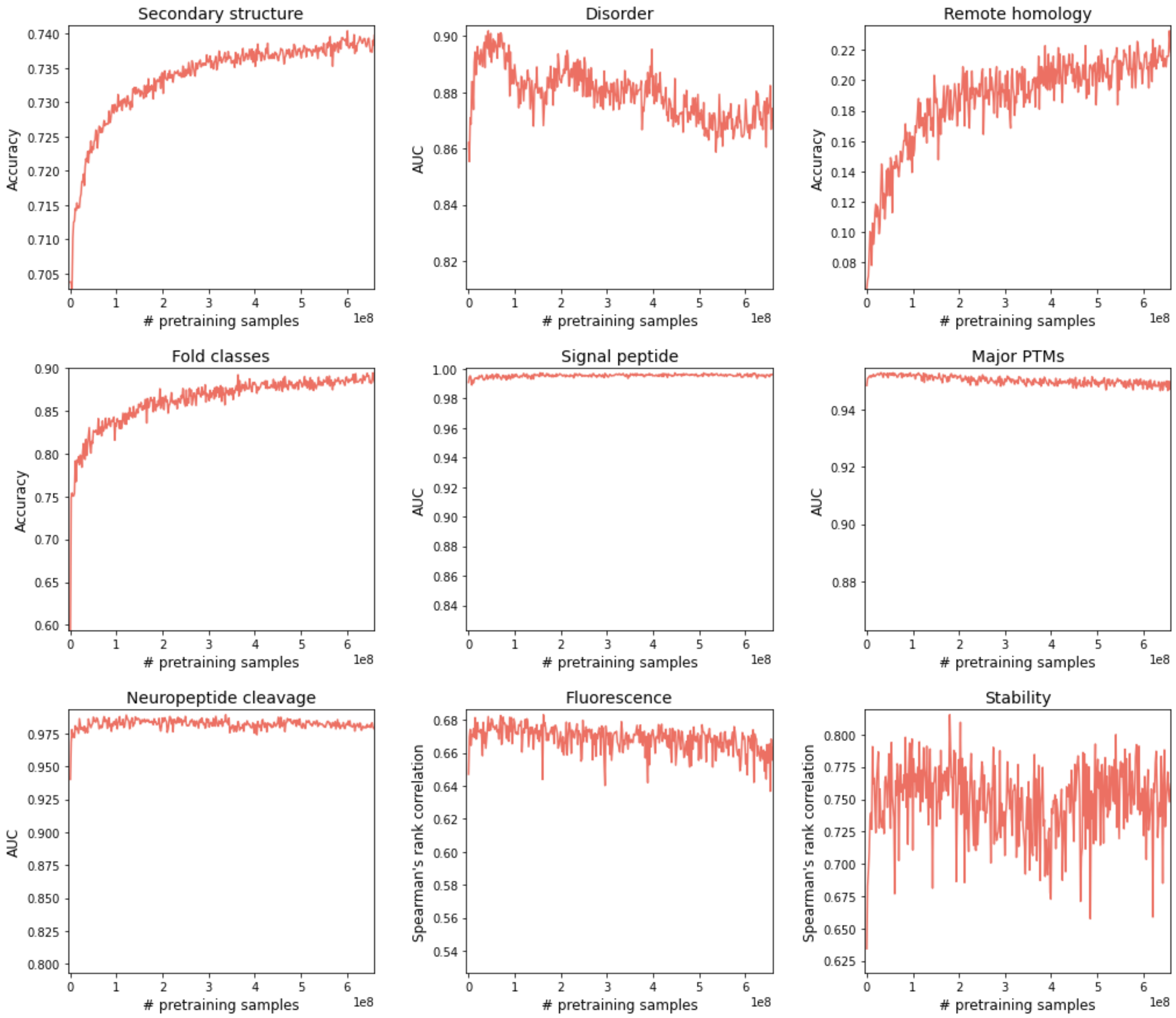


**Supplementary Fig. S1: The impact of pretraining across all nine benchmarks**

Performance of fine-tuned ProteinBERT models over the 9 tested benchmarks (Table 1) as a function of pretraining amount (measured by the number of processed proteins).

### Supplementary Methods

#### Determining the GO annotations associated with each UniRef90 record

Metadata on all possible GO annotations (47,376 annotations) was downloaded from CAFA [1] at https://www.biofunctionprediction.org/cafa-targets/cafa4ontologies.zip. This metadata included information about the tree structure of GO (i.e. the parent terms that each term is descended from). From UniRef90, we extracted the GO annotations associated with each protein (by extracting all properties of type “GO Molecular Function”, “GO Biological Process” or “GO Cellular Component”). For each GO annotation reported for a given protein, we also included all its parent annotations (up to the root nodes of the GO tree). We then filtered only the 8,943 of the 47,376 GO terms that occurred at least 100 times among the ~106M parsed UniRef90 records. The code for processing UniRef90 and its associated GO annotations is in the *create_uniref_db* script on the project’s GitHub repository (https://github.com/nadavbra/protein_bert).

#### Benchmarks

We used 9 benchmarks overall: 4 from TAPE [2] (left unchanged), and 5 additional ones derived from a variety of sources. All 9 benchmark datasets are available via FTP at ftp.cs.huji.ac.il/users/nadavb/protein_bert/protein_benchmarks.

The 4 TAPE benchmarks are: i) secondary structure prediction at the residue level, out of 3 classes (Helix, Strand and Other); ii) remote homology detection as a multiclass prediction of structural fold, out of 1,195 families [3, 4]; iii) fluorescence prediction for mutated protein sequences [5]; and iv) intrinsic protein stability [6]. We also created an additional benchmark of disorder prediction, derived from TAPE’s secondary structure benchmark [7].

The Major PTMs benchmark, whose task is predicting whether each residue undergoes any form of post-translational modification, was acquired from PhosphoSitePlus [8]. Specifically, we used “PhosphoSite seq”, a dataset provided by PhosphoSitePlus where all residues known to be post-translationally modified are labeled.

The signal peptide benchmark, whose task is predicting whether an entire protein sequence has a signal peptide or not, was derived from signalP 5.0 [9] by combining the dataset across all kingdoms.

The neuropeptide cleavage benchmark was taken from an earlier work [10, 11]. Here, the goal is to predict if a basic residue (K or R) undergoes cleavage, where all candidate sequences have a signal peptide.

In the fold classes benchmark the goal is to predict the high-level structural class of each protein, out of the 7 SCOP (Structural Classification of Proteins) classes. Sequences were filtered according to Astral SCOPe 2.07 genetic domain sequence subsets, with less than 40% identity to each other [3, 4].

For all TAPE datasets we used the predefined training, evaluation and test partitions (reporting results on the test sets). For the other benchmarks (which we derived ourselves), we randomly split them into training, evaluation and test sets.

### Pretraining episodes

Pretraining was performed in episodes of homogenous encoding sequence lengths: 128, 512 or 1,024 tokens. The ~106M records in the UniRef90 dataset were randomly shuffled and read into memory in chunks of 100,000 records. Each protein record was randomly assigned to one of the three types of episodes in a way that favored proteins to be assigned to an episode of sequence length similar to the length of the protein. Specifically, let $L_0$ be the length of a candidate protein and let $L_1 \in \{128, 512, 1024\}$ be the encoding sequence length of an episode. Defining $r = \max\left\{\frac{L_0}{L_1}, \frac{L_1}{L_0}\right\} \geq 1$ to be the ratio of these two lengths, we made the probability of assigning the candidate protein to that episode to be proportional to $\exp(-r)$.

Every 15 minutes of pretraining, the training episode was changed into the episode with the largest number of pending records (with an option to continue the same episode, if it still had the highest number of pending records). A new chunk of 100,000 records was read whenever a running episode was left without any pending records, or when no pending records were left at all. The code for managing the episodes is available as part of the *pretrain_proteinbert* script on the project's GitHub repository (https://github.com/nadavbra/protein_bert).

### Optimization

Pretraining of ProteinBERT on protein sequences and GO annotations derived from UniRef90 was carried out with a LAMB optimizer [12] with an initial learning rate of 2e-04. Learning rate was then reduced on plateau using Keras's *ReduceLROnPlateau* [13] (with parameters *patience = 20, factor = 0.8, min_lr = 5e-05*). Fine-tuning of ProteinBERT on downstream tasks was then done with an ADAM optimizer [14]. For the first epochs, when all layers except the final fully-connected layer were frozen, we used an initial learning rate of 1e-02. For the next epochs allowing fine-tuning of all model parameters we used an initial learning rate of 1e-04, and for the final epoch (with a larger sequence length) we used a learning rate of 1e-05. For the fine-tuning as well we used Keras's *ReduceLROnPlateau* (see next section for parameter details). During pretraining we used batch sizes of 128, 64 or 32 for episodes of 128, 512 or 1,024 tokens, respectively. During fine-tuning we used a batch size of 32.

### Fine-tuning and evaluation

When fine-tuning ProteinBERT on a downstream task with a labeled dataset (one of the nine benchmarks listed in Table 1), a new fully-connected layer was added to its output. For tasks involving local predictions (at the residue level), the final fully-connected layer was connected to its local representations (Fig. 1). Specifically, we concatenated all the local normalization layers (2 per transformer block) and the output sequence, and connected them to the final fully-connected layer. Likewise, for tasks involving global predictions (at the entire protein level), the final layer was connected to its global representations. Specifically, we used a concatenation of all the global normalization layers (2 per transformer block) and the output annotations. In all cases we used dropout with a rate of 0.5 for the concatenation of these hidden layers (which was connected to the final fully-connected layer), but kept all other layers unregularized.

For each dataset we derived separate training, validation and test sets. For benchmarks without a validation set, we randomly split 10% of the training set into an independent validation set through stratified sampling (using scikit-learn's *train_test_split* [15]), and kept only the other 90% for training.

Throughout the entire fine-tuning process we used a sequence length of 512 tokens, except for a final epoch of 1,024 tokens which was introduced to encourage the model to generalize to different sequence lengths. For all but the last epoch, we filtered out all the proteins larger than 512 tokens from the training and validation set, and trained the model on the remaining records. For the final epoch, we trained the model on records from the training and validation set of size up to 1,024 tokens.

By measuring the loss on the validation set at the end of each epoch, we applied early stopping using Keras's *EarlyStopping* [13] (with parameters *patience = 2, restore_best_weights = True*). We also reduced the learning rate on plateau using Keras's *ReduceLROnPlateau* (with parameters *patience = 1, factor = 0.25, min_lr = 1e-05*).

Evaluation of the fine-tuned model was carried over the held-out test set. We applied the model on all test-set records by starting with a sequence length of 512 tokens and increasing the length by factors of 2 each time, such that each test-set protein was processed with the minimal sequence length of power of 2 that is larger than the length of that protein (e.g. a protein of 80 tokens would be processed with a length of 512, a protein of 800 tokens would be processed with a length of 1,024, and a protein of 10,000 tokens would be processed with a length of 16,384). Notably, even though fine-tuning was carried over proteins of up to 1,024 tokens (with all but the final epoch being restricted to 512 tokens), ProteinBERT still generalized reasonably well to longer sequences (Fig. 4).

The code for fine-tuning and evaluating ProteinBERT on downstream tasks is available on the project's GitHub repository (https://github.com/nadavbra/protein_bert).

### Experimenting with other tokenizations

In its current form, ProteinBERT uses individual amino acids as tokens [16]. During its development, we also experimented with wordpiece and sentencepiece, namely using n-grams of amino acids as tokens (instead of individual residues). However, the large vocabulary size led to increased memory and runtime requirements, but not to improved performance. In some fine-tuning tasks, we even saw reduced performance. Therefore, we eventually decided to abandon this approach and settled on simple, character-level tokenization.

### References


1. Zhou N, Jiang Y, Bergquist TR, et al (2019) The CAFA challenge reports improved protein function prediction and new functional annotations for hundreds of genes through experimental screens. Genome Biol 20:1–23
2. Rao R, Bhattacharya N, Thomas N, et al (2019) Evaluating protein transfer learning with tape. Adv Neural Inf Process Syst 32:9689
3. Andreeva A, Howorth D, Chothia C, et al (2014) SCOP2 prototype: a new approach to protein structure mining. Nucleic Acids Res 42:D310-4. https://doi.org/10.1093/nar/gkt1242
4. Andreeva A, Kulesha E, Gough J, Murzin AG (2020) The SCOP database in 2020: expanded classification of representative family and superfamily domains of known protein structures. Nucleic Acids Res 48:D376--D382
5. Sarkisyan KS, Bolotin DA, Meer M V, et al (2016) Local fitness landscape of the green fluorescent protein. Nature 533:397–401
6. Rocklin GJ, Chidyausiku TM, Goreshnik I, et al (2017) Global analysis of protein folding using massively parallel design, synthesis, and testing. Science (80- ) 357:168–175
7. Moult J, Fidelis K, Kryshtafovych A, et al (2018) Critical assessment of methods of protein structure prediction (CASP)—Round XII. Proteins Struct Funct Bioinforma 86:7–15
8. Hornbeck P V, Zhang B, Murray B, et al (2015) PhosphoSitePlus, 2014: mutations, PTMs and recalibrations. Nucleic Acids Res 43:D512--D520
9. Armenteros JJA, Tsirigos KD, Sønderby CK, et al (2019) SignalP 5.0 improves signal peptide predictions using deep neural networks. Nat Biotechnol 37:420–423
10. Ofer D, Linial M (2015) ProFET: Feature engineering captures high-level protein functions. Bioinformatics 31:3429–3436
11. Brandes N, Ofer D, Linial M (2016) ASAP: A machine learning framework for local protein properties. Database 2016:. https://doi.org/10.1093/database/baw133
12. You Y, Li J, Reddi S, et al (2019) Large batch optimization for deep learning: Training bert in 76 minutes. arXiv Prepr arXiv190400962
13. Chollet F, others (2015) keras
14. Kingma DP, Ba J (2014) Adam: A method for stochastic optimization. arXiv Prepr arXiv14126980
15. Pedregosa F, Varoquaux G, Gramfort A, et al (2011) Scikit-learn: Machine learning in Python. J Mach Learn Res 12:2825–2830
16. Ofer D, Brandes N, Linial M (2021) The language of proteins: NLP, machine learning \& protein sequences. Comput Struct Biotechnol J
